# Proposing a Systematic Lineage Classification Below the Genotype Level for Dengue Serotypes 1 and 2

**DOI:** 10.1101/2024.03.25.586629

**Authors:** James Siqueira Pereira, Gabriela Ribeiro, Vinicius Carius de Souza, Isabela Carvalho Brcko, Igor Santana Ribeiro, Iago Trezena Tavares De Lima, Svetoslav Nanev Slavov, Sandra Coccuzzo Sampaio Vessoni, Maria Carolina Elias, Alex Ranieri Jerônimo Lima

## Abstract

Dengue virus (DENV), a mosquito-borne flavivirus, is causing a significant outbreak in Brazil. The recent surge in complete DENV genome sequences necessitates a standardized classification system for an improved understanding of viral dynamics and transmission patterns. Traditionally, DENV classification relies on serotypes and genotypes but lacks a consensus for sub-genotype classification. This hinders comprehensive analyses of viral diversity. We address this gap by proposing a novel lineage classification system for DENV using a semi-automatic workflow, leveraging the use of complete genome sequences to classify and re-evaluate DENV genetic diversity. This system offers a more granular classification scheme compared to current methods. The proposed hierarchical nomenclature, incorporating serotype, genotype, subgenotype, lineage, and sublineage, facilitates precise tracking of viral introductions and evolutionary events. This information might have crucial implications for public health interventions, enabling more targeted control strategies and improved monitoring of vaccine effectiveness during future outbreaks.

## Introduction

Dengue fever, caused by the four Dengue virus (DENV) serotypes, stands as a paramount arboviral ailment globally. Manifestations range from mild febrile illness to severe hemorrhagic diathesis, also recognized as Dengue Shock Syndrome (Wang *et al*., 2020). Transmitted by *Aedes* mosquito vectors, notably *Aedes aegypti* and *Aedes albopictus*, the disease exhibits an expanding geographical footprint, portending an inevitable surge in DENV-related morbidity and mortality rates worldwide. Notably, in 2023, the global toll encompassed over five million reported cases of dengue, accompanied by more than 5000 fatalities across 80 countries. The Americas Region bore the brunt, recording approximately 4.1 million cases, representing nearly 80% of the global burden (WHO, 2023). This high transmission rate has continued into 2024, with a total of 1,874,021 suspected cases of dengue reported spanning from epidemiological weeks 1 to 8. Out of these cases, 658,215 were lab-confirmed, 1,670 were classified as severe (0.1%), with 422 resulting in fatalities (case fatality rate 0.023%) (PAHO, 2024).

DENV belongs to the *Flaviviridae* family with a well-defined genomic structure: a single positive strand RNA of approximately 11 Kb. The genomic organization encompasses two untranslated regions (UTR) at the 5’ and 3’ ends, flanking an Open Read Frame (ORF), which serves as a template to encode a polyprotein composed of 3 structural proteins and 7 non-structural proteins. The structural proteins are the capsid (C), pre-membrane (PrM), and envelope (E), while the non-structural proteins are numbered as NS1, NS2A, NS2B, NS3, NS4A, NS4B, and NS5 (Nanaware *et al*., 2021). There are four distinct DENV serotypes (DENV-1-4), along with a recently described fifth serotype (DENV-5) (Mustafa *et al*., 2015). In addition to this classification, each serotype is divided into different genotypes, often defined as strains with less than 6% divergence at the junction between the E and NS1 coding regions (Rico-Hesse, 1990).

The elucidation of virus lineages and strains is pivotal for comprehending their evolutionary trajectories. Integrating this genetic information with diverse datasets, including geographical, epidemiological, and clinical parameters, furnishes invaluable insights into the disease dynamics (Grubaugh, 2022). Previous studies suggest that DENV lineages go extinct every 7-10 years, being replaced by new lineages, typically resulting in epidemics (Adams *et al*., 2006; Nunes *et al*., 2014; Lourenço *et al*., 2018).

Nonetheless, the absence of a standardized lineage classification and nomenclature for DENV at subgenotype levels poses a significant challenge in longitudinally monitoring lineage substitutions. Moreover, communicating lineage-replacement events or identifying epidemic strains to stakeholders and public health agencies becomes problematic in the absence of an established lineage system. The utilization of the lineage classification system proposed by Rambaut et al., 2020 has successfully overcome those issues during the COVID-19 pandemic, as it helped to precisely track the virus variants across the globe for SARS-CoV-2 (Oude Munnink and Koopmans, 2023; WHO, 2024). This approach has since been adapted to propose lineage designation frameworks for other viruses, including Monkeypox, Rabies, and HRSV (Campbell *et al*., 2022; Happi *et al*., 2022; Goya *et al*., 2024), leveraging the rationale behind its implementation.

In March 2022, the World Health Organization (WHO) launched the Global Arbovirus Initiative, a collaborative effort designed to unite key stakeholders in bolstering surveillance and preventative strategies against arboviruses (Balakrishnan, 2022). Leveraging existing genomic surveillance networks established for SARS-CoV-2, this initiative facilitated the real-time availability of comprehensive DENV genomes. With the escalation of the number of dengue cases came a pressing necessity to systematically organize and centralize this burgeoning genomic data, culminating in establishing the GISAID EpiArbo database in 2023 (Wallau *et al*., 2023). Concurrently, novel bioinformatic tools emerged, geared towards impartially proposing and delineating new lineage classification systems for pathogens, adept at managing vast genomic datasets (Campbell *et al*., 2022; Ha and Aylward, 2024; McBroome *et al*., 2024)

Brazil has become a significant area of concern due to the rapid rise in dengue cases. During the first five epidemiological weeks of 2024, a substantial number of 455,525 dengue cases were reported in Brazil (PAHO and WHO, 2024). This concerning trend continued, with an additional 239,058 cases reported by epidemiological week eight (PAHO, 2024). Notably, while all four DENV serotypes have been identified in Brazil, in recent years there is also intensive circulation of DENV serotypes 1 and 2 (Figure S1). This regional dominance underscores the urgent need for tailored strategies aimed at classification and surveillance to manage and control dengue within the country’s borders effectively. As such, understanding the dynamics of these prevalent serotypes beyond the genotype level within Brazil holds significant implications for global dengue control efforts, necessitating a concerted focus on this geographic hotspot. Thus, we propose a lineage classification system for DENV serotypes 1 and 2 in this work, applying a semi-automatic workflow.

## Methods

### Retrieving and filtering the sequence dataset

The genomes of DENV-1 and DENV-2 were obtained from GISAID EpiArbo (https://www.gisaid.org/) selecting sequences with a minimum 10,000-nt, aiming to retrieve only genomes expected to be complete. Only DENV original passages and genomes obtained from the human host were selected. The DENV-1 and DENV-2 datasets contained sequences obtained up to January 2024 with initial collection dates between 1944 and 1976. To ensure comprehensive data coverage, we employed an in-house established script to assess the presence of ambiguous bases in each genome, setting a 10% cutoff. This strategy ensured that only genomes with at least 90% coverage would be retained. The metadata of the sequences used (including the GISAID accession IDs) are available in Supplementary files 1 and 2.

### Phylogenetic tree reconstruction

Genomic sequences from the two filtered serotype datasets were aligned independently using the Augur align tool within the Augur program toolkit v24.1.0 (Huddleston *et al*., 2021), employing the MAFFT v.7.520 algorithm (Katoh and Standley, 2013). Subsequently, maximum likelihood (ML) phylogenetic trees were constructed utilizing IQ-TREE v2.2.6 (Minh *et al*., 2020). The trees were reconstructed employing the GTR+F+I+G4 nucleotide substitution model, selected by the ModelFinder application (Kalyaanamoorthy *et al*., 2017) within IQ-TREE2. To assess the robustness of tree topology, 5000 UFBoot replicates were generated. Phylogenetic trees were dated using Treetime v0.11.2 (Sagulenko, Puller and Neher, 2018), integrated in Augur. Furthermore, multiple refinements were applied during this phase, including rerooting the tree using the midpoint method, inference of node confidence dates, stochastic resolution of polytomies, and pruning of genomes that exhibited deviation from the evolutionary rate (clock filter interquartile distance of 4). To automate the analysis workflow, a local Bash script was developed and is available on the GitHub repository (https://github.com/alex-ranieri/denvLineages/blob/main/scripts/execPhyloDenv.sh).

Ancestral inference and identification of nucleotide mutations were conducted using Augur ancestral, while amino acid mutations were annotated utilizing Augur translate. For DENV-1, the GenBank sequences NC_001477.1 served as the reference, and for DENV-2, NC_001474.2 was utilized.

### Lineage definition workflow

Initially, DENV-1 and DENV-2 lineages were defined using an automated heuristic approach implemented in Autolin (McBroome *et al*., 2024). The code was modified to consider only clusters with robust statistical support (UFBoot ≥ 90%) (available at https://github.com/alex-ranieri/denvLineages/blob/main/scripts/annotate_json.py). This lineage definition method strictly considered the occurrence of, at least, one amino acid mutation within minimally sized groups (n ≥ 10) where at least 90% of sequences presented the mutation at every possible hierarchical depth level. To define a lineage, it was considered a GRI (Genotype Representation Index) of 1, implying, on average, that a single amino acid mutation differentiates the lineage from a randomly chosen sample on the phylogenetic tree.

The Autolin output underwent post-processing using a custom Python script (available at https://github.com/alex-ranieri/denvLineages/blob/main/scripts/posProcessingAutolin_aa.py) to ensure alignment with Augur Clades standards, as implemented within the Augur v24.1.0 toolkit, which became the primary platform for lineage assignment. This post-processing step involved manual curation of both the lineage designations generated by Autolin and the corresponding mutation table. The curation process focused on retaining only those mutations observed in all, or at least 90%, of sequences within a given branch. This selection strategy aimed to identify mutations with high discriminatory power, capable of reliably differentiating the corresponding phylogenetic nodes.

To ensure lineage assignments transcended serotype boundaries and adhered to established classification criteria, a systematic approach was implemented to link lineage-defining mutations with those previously associated with distinct genotypes.

This validation process leveraged a hereditary framework, where mutations associated with designated lineages were compared against the established genotype mutation table curated within the Nextstrain dengue build (https://github.com/nextstrain/dengue). This methodology was consistently applied across all hierarchical levels, encompassing both primary lineages and any identified sublineages.

### Lineage labeling definitions

The proposed nomenclature system for DENV lineages follows a hierarchical structure of serotype, genotype, subgenotype, lineage, and sublineage, adapting the rationale used in the Pango system (Rambaut *et al*., 2020). At the genotype level, a combination of numbers and letters designated both the serotype and genotype classification. For instance, “1V” means DENV-1 genotype V. The subgenotype nomenclature was chosen because it represents the initial hierarchical subdivision designated by Autolin software, following the genotype level. Thus, subgenotypes were defined using a period (“.”) followed by a capital letter (“A,” “B,” etc.) assigned sequentially. This alphabetical attribution commenced with the letter “A” for clarity and ease of reference. For instance, “1IV.A” represents a subgenotype of DENV-1 genotype IV. At the same time “1V.A” denotes a subgenotype of DENV-1 genotype V. Lineages and sublineages were labeled using ascending numeric identifiers separated by periods to indicate hierarchical depth. For clarity, this labeling system is limited to three numerical levels. When the fourth level is reached, an additional letter followed by a period and another number is appended to the lineage label. Following this logic, “2II.D.B.1” signifies a sublineage derived from lineage “2II.D.10.1.2” instead of using a longer string like “2II.D.10.1.2.1”. The labeling scheme reflects the order of nodes within each genotype based on the tree topology.

### Implementation of lineage assignment

To facilitate the classification of novel DENV sequences into defined lineages, a dedicated Nextclade dataset was constructed. This dataset (available at https://github.com/alex-ranieri/denvLineages/tree/main/Nextclade_V2_data) serves as a reference for lineage assignment and leverages the Nextclade framework’s robust phylogenetic inference capabilities (Aksamentov *et al*., 2021). By incorporating this dataset within the lineage assignment workflow, newly obtained DENV sequences can be efficiently classified based on their genetic relatedness to previously defined lineages.

## Results

### Phylogenetic lineage-labeled tree

A total of 5,084 and 3,672 DENV-1 and DENV-2 genomes, respectively, were employed to construct the lineage-labeled phylogenetic trees. The temporal distribution of these sequences across collection years is depicted in Figure 1. Furthermore, the geographical distribution across continents and countries is presented in Figure S2. Autolin successfully assigned lineage labels to most of the DENV-1 and DENV-2 sequences. It labeled 5,036 (DENV-1) and 3,668 (DENV-2) sequences, representing approximately 99% of each dataset. This translates to a high level of representativeness for the lineage assignment process. For DENV-1, Autolin generated 192 unique annotation labels across 6 hierarchical levels, while for DENV-2, 148 unique labels were generated across 8 hierarchical levels. Details regarding the GRI values and initial labels are provided in Supplementary Files 3 and 4. The top hierarchical levels annotated by Autolin (i.e., the inner nodes of the tree) reflected the DENV genotype labels.

**Figure 1.**
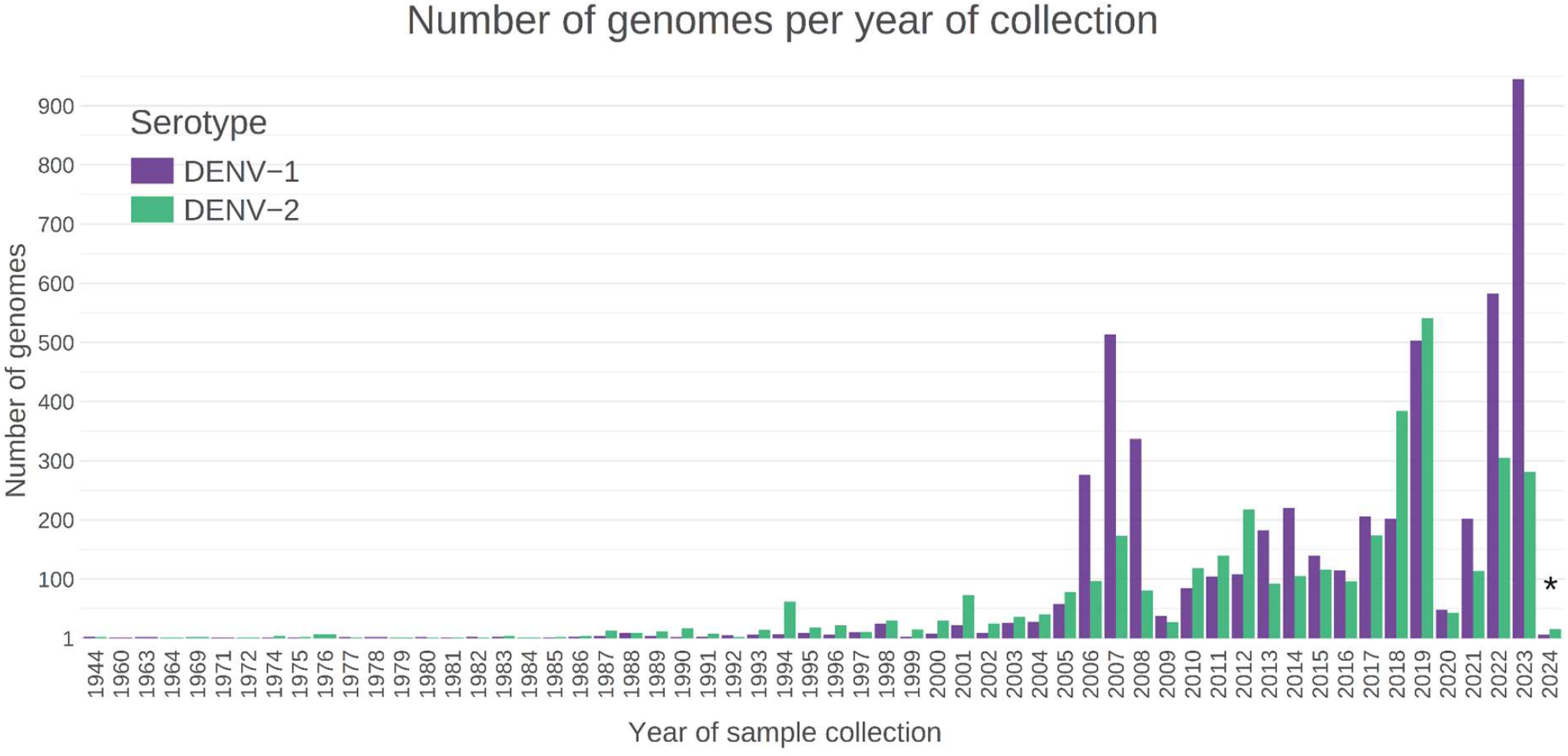
Temporal Distribution of DENV Genomes Employed for Lineage Assignment. The data are grouped by collection date. * For 2024, sequences were retrieved from GISAID up to January 4th for DENV-1 and January 11th for DENV-2.

Following manual curation and application of the established lineage labeling definitions, a comprehensive DENV lineage classification system was developed (Figure 2). Notably, for DENV-1, a novel genotype containing 22 sequences (named 1VI) was identified, which was observed circulating in Africa with the most recent sample collected in 2021, and with no assigned subgenotypes. In total, 26 subgenotypes were designated for DENV-1, encompassing 86 distinct lineages and their sublineages. Interestingly, subgenotype designations were absent for 1II, while 1V displayed the highest subgenotype diversity within DENV-1 (n=20).

**Figure 2.**
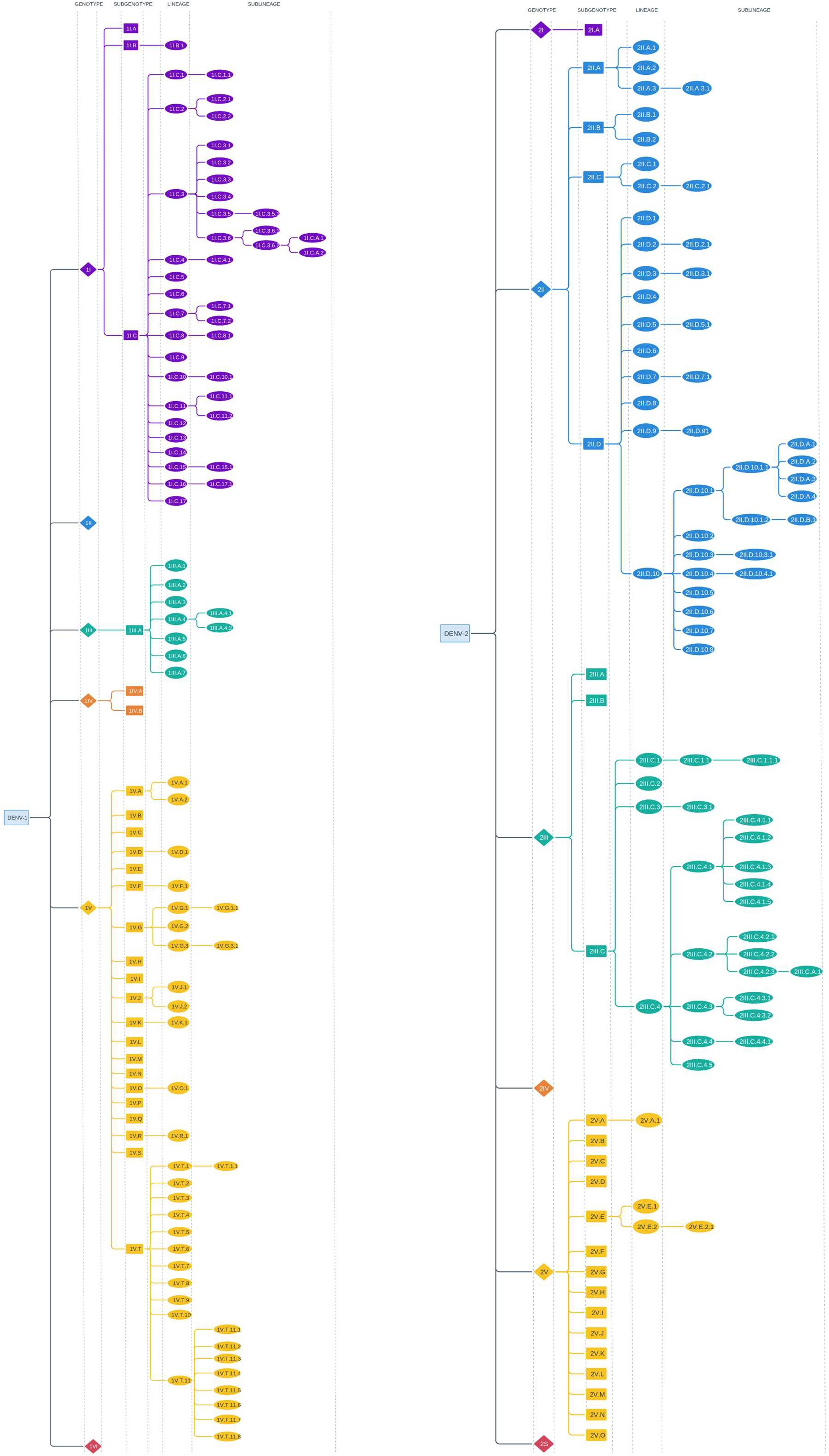
DENV Lineage Classification System. Schematic depicting the proposed DENV lineage classification system for serotypes 1 (left) and 2 (right). The system incorporates a hierarchical structure for each serotype, encompassing four main levels displayed from left to right: genotype, subgenotype, lineage, and sublineage.

For DENV-2, 23 subgenotypes were designated, encompassing 66 lineages and their sublineages. The genotype 2V exhibited the greatest subgenotype richness within DENV-2 (n=15), whereas 2IV lacked subgenotype designations. Lineage-labeled phylogenetic trees for DENV-1 and DENV-2 are available for visualization at https://nextstrain.org/fetch/raw.githubusercontent.com/alex-ranieri/denvLineages/main/Nextclade_V2_data/DENV1/tree.json?branchLabel=lineage&c=lineage_membership&p=full and https://nextstrain.org/fetch/raw.githubusercontent.com/alex-ranieri/denvLineages/main/Nextclade_V2_data/DENV2/tree.json?branchLabel=lineage&c=lineage_membership&p=full, respectively.

In DENV-1 lineage classification, the NS5, E, and NS3 genes served as the primary determinants, being utilized in 98 (34.5%), 44 (15.5%), and 35 (12.3%) instances, respectively. Similarly, DENV-2 lineage definitions were primarily established based on mutations within the NS5, E, and NS2A genes, with these genes being employed in 64 (29.1%), 35 (15.9%), and 31 (14.1%) instances, respectively. A comprehensive mutation table, compatible with the Augur Clades, has been compiled and is available for reference at https://github.com/alex-ranieri/denvLineages/tree/main/mutation_tables.

### Implementation of the lineage classification system

To promote the widespread adoption of the proposed DENV lineage classification system, a dedicated Nextclade dataset was constructed. A web version for DENV-1 and DENV-2 are available for lineage assignment at https://v2.clades.nextstrain.org/?dataset-url=https://github.com/alex-ranieri/denvLineages/tree/main/Nextclade_V2_data/DENV1 and https://v2.clades.nextstrain.org/?dataset-url=https://github.com/alex-ranieri/denvLineages/tree/main/Nextclade_V2_data/DENV2, respectively. This dataset serves as a reference resource for assigning the lineage of novel DENV genomes generated by ongoing genomic surveillance initiatives. To evaluate the accuracy of the system in assigning lineages to novel sequences, a validation approach was employed using Nextclade v2.14.0. This involved the use of genomic sequences (complete or partial) deposited in GISAID during 2024. Forty DENV-1 sequences with collection dates ranging from January 5th to 31st and 14 DENV-2 sequences collected between January 12th and 30th were selected for analysis (data accessed on GISAID on March 20th, 2024). All obtained samples were from Brazil.

Analysis of DENV-1 sequences revealed their assignment to subgenotypes 1V.T (n=31) and 1V.J (n=8), encompassing their constituent lineages. Notably, one sequence was classified solely to genotype 1V without subgenotype designation (Figure 3 left). The genomic coverage of these sequences ranged from 74% to 97%. Both subgenotypes were first identified in 2021 and have exhibited ongoing circulation (including their associated lineages) within Brazil up to the beginning of 2024. Analysis of DENV-2 sequences revealed that a majority of sequences (n=12) were assigned to lineage 2II.D.10.1.1 (Figure 3 right). This lineage was first detected in Peru in September 2019, but its subsequent circulation has been predominantly observed in Brazil. Notably, additional detections of 2II.D.10.1.1 were identified in Paraguay and Bolivia. The most recent samples for this lineage were collected in Brazil during 2024, suggesting ongoing circulation. Lineage 2III.C.4.1 was primarily identified in Brazil, being first detected in 2016, with the most recent sequence collected in August 2023. Finally, one sequence was classified solely to genotype 2V (Asian I) without further lineage designation. The genomic coverage of DENV-2 genome sequences ranged from 74% to 99%.

**Figure 3.**
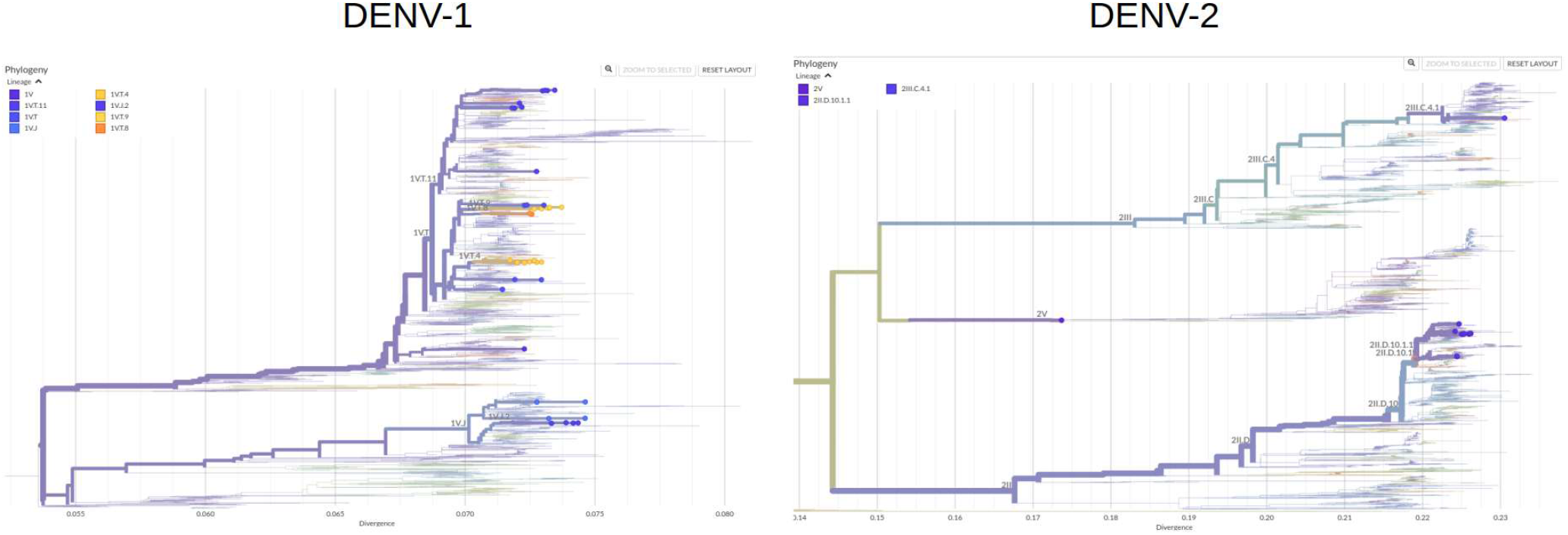
Genetic Diversity of Dengue Virus (DENV) Sequences in Brazil, 2024. Analysis of distinct subgenotype distributions for DENV-1 (40 sequences) is on the left panel; DENV-2 (15 sequences) sequences are displayed on the right panel. GISAID accession IDs can be found in Supplementary Files 5 and 6.

## Discussion

The escalating arboviral threat posed by DENV necessitates the development of a robust, global genotyping framework. This framework would facilitate the contextualization of spatiotemporal epidemiological data, enabling a deeper understanding of DENV dynamics. Endemic persistence and incursions into previously non-endemic regions have significantly shaped DENV’s genetic diversity and evolutionary divergence (Chen and Vasilakis, 2011; Fontaine *et al*., 2018).

DENV exhibits significant genetic heterogeneity, organized hierarchically into serotypes, genotypes, and subgenotype clades. However, inconsistent nomenclature systems, a lack of standardized classification procedures, and ambiguities in reporting antigenic diversity hinder comprehensive analysis (Cuypers *et al*., 2018). A recent study proposed a unified global genotyping framework for DENV-1, utilizing complete E gene sequences from diverse epidemic regions to address these concerns (Li *et al*., 2022). This framework offers a standardized approach, revealing distinct global epidemic patterns, identifying persistent transmission zones and emerging outbreaks, and highlighting the necessity for a coordinated global surveillance platform to combat DENV’s expansion. However, there is a lack of lineage classification systems implemented within bioinformatic tools for DENV classification below the genotype level.

Our study addresses this gap by proposing a novel nomenclature system for DENV lineages, currently encompassing DENV-1 and DENV-2. This system aims to enhance viral surveillance and classification addressing key aspects. Firstly, it will facilitate improved monitoring efforts by providing a clear and consistent method for lineage assignment. Secondly, it will enable researchers to more precisely track the introduction and dissemination of specific regional lineages. Most importantly, our system acknowledges the growing volume of complete DENV genomes sequenced in recent years, promoting improved identification, reporting, and understanding of the virus.

DENV-1, Brazil’s most prevalent and continuously circulating serotype, is traditionally classified into five genotypes (Goncalvez *et al*., 2002). To enhance the resolution of DENV-1 classification, we propose a more granular system that categorizes this serotype into 26 subgenotypes and 86 lineages. This refined approach facilitates more precise identification and reporting of analyzed genomic samples. For instance, a study by Adelino et al., 2021, sought to characterize the diversity of DENV-1 genotype V, identifying three distinct clades (I, II, and III). The authors suggested that clades II and III replaced clade I in 2019. When analyzed using the proposed classification system, the sequences from those clades were grouped as follows: Clade I - Lineage 1V.A.1; Clade II - 1V, 1V.G.1 and 1V.G.1.1; Clade III - 1V. This finding shows that a concise lineage classification system allows for precise reporting of replacement events. It would be more informative to say that lineages 1V.G.1 and 1V.G.1.1, along with a diverging 1V branch, replaced lineage 1V.A.1. Moreover, an additional evolutionary event was observed within Clade II with the advent of the sublineage 1V.G.1.1, allowing precise tracking of genetic diversity. Finally, it shows that the detected replacement events occurred not necessarily spanning new lineages.

Intriguingly, the proposed classification system identified a putative sixth genotype for DENV-1, labeled as 1VI, and covers 22 sequences. These sequences are currently classified as part of genotype V based on the mutation scheme used by the Nextstrain dengue compilation (https://github.com/nextstrain/dengue). This discovery demonstrates the potential of the proposed system to uncover previously unknown diversity within the DENV-1. The 1VI genotype has been circulating in Africa, and the most recent sample collected in 2021 does not have assigned subgenotypes. Identifying such potentially divergent lineages is crucial for evolutionary and epidemiological monitoring, particularly for tracking the emergence and re-emergence of strains in regions with limited genomic surveillance capabilities.

In investigations into the introduction of DENV-2 in Brazil, it was also observed that replacement events occurred within genotype III (Asian/American) of DENV-2 with the introduction of a fourth lineage. In a study conducted by De Jesus et al., 2020, the authors identified a sequence cluster naming it as Clade BR-4. Our classification method found that this cluster of sequences belongs to the same subgenotype (2III.C), specifically to the 2III.C.4 lineage. However, they could be distinguished into two different sublineages, with 17 sequences classified as 2III.C.4.1 and two classified as 2III.C.4.2. It is worth noting that 11 of the 17 sequences assigned to 2III.C.4.1 were classified into a sublineage called 2IIIC.4.1.4, adding a deeper comprehension on the cluster evolutionary dynamics.

As performed in the Clade BR-4 analysis, we applied our classification system to the samples from Amorim et al., 2024, which provides valuable insights into the genetic diversity of DENV-2 and its implications for epidemiology. In that study, the authors identified a group of samples calling it Lineage 5, belonging to genotype II (Cosmopolitan). These samples were cohesively grouped in our proposed classification as subgenotype 2II.D, specifically to sublineage 2II.D.10.1 within lineage 2II.D.10. However, two sequences present in Lineage 5 reported by Amorim et al., 2024, could be classified at a further level of depth, belonging to sublineage 2II.D.10.1.1, indicating genetic diversification events within the previously reported Lineage 5.

By investigating the genetic diversity of Clade BR-4 and Lineage 5, we inferred that each of these sequence groups is indeed internally related, as indicated by the authors, as they share the ancestries of lineages 2III.C.4 and 2II.D.10.1, respectively. However, they exhibit internal diversification events that should be considered from a genomic surveillance perspective.

The proposed unified classification system for DENV represents a significant advancement over scattered generic nomenclatures. This detailed and precise approach not only facilitates dynamic DENV monitoring, crucial for targeted vaccination strategies during outbreaks and vaccine introductions but also enables a more agile response to changes in the epidemiological landscape. By allowing for specific lineage and sublineage monitoring, the system simplifies communication with public health authorities and facilitates effective tracking of potential viral replacements. Additionally, this standardized tool simplifies the interpretation of phylogenetic analyses across studies, resulting in a more accurate understanding of DENV evolution and dispersal, ultimately informing the development of effective control and prevention strategies.

### Future perspectives

This study establishes the foundation for applying the proposed Dengue virus (DENV) lineage classification system to DENV-3 and DENV-4. This expansion will create a comprehensive framework for DENV lineage designation encompassing all four serotypes.

The system’s user-friendliness is augmented by its online accessibility through the Nextclade V2 web application for DENV-1 (https://v2.clades.nextstrain.org/?dataset-url=https://github.com/alex-ranieri/denvLineages/tree/main/Nextclade_V2_data/DENV1) and DENV-2 (https://v2.clades.nextstrain.org/?dataset-url=https://github.com/alex-ranieri/denvLineages/tree/main/Nextclade_V2_data/DENV2). Alternatively, users can download datasets from https://github.com/alex-ranieri/denvLineages/tree/main for offline analysis using Nextclade CLI V2.14.0. Future integration with the Viral Identification Pipeline for Emergency Response (VIPER) assembly pipeline (https://github.com/alex-ranieri/viper) is planned. Additionally, datasets will be upgraded to ensure compatibility with Nextclade V3.

To facilitate the proposition of novel lineages, a comprehensive guideline document will be developed and hosted on https://github.com/alex-ranieri/denvLineages/tree/main. Finally, we aim to create a dedicated pipeline that leverages Nextclade, this DENV lineage classification system’s datasets, and Autolin software to suggest new lineages for newly submitted sequences. This will further enhance DENV genomic surveillance efforts.

## Acknowledgements

Funding for this work was provided by Fundação Butantan and the São Paulo Research Foundation (FAPESP) through grant number 21/11944-6, titled “Continuous improvement of vaccines: Center for Viral Surveillance and Serological Assessment (CeVIVAS)”. We express our sincere gratitude to all researchers who deposited their sequences on GISAID, particularly those affiliated with the Central Public Health Laboratories (LACEN) from the Brazilian States of Alagoas, Pará, and Paraná. We are also grateful to São Paulo City Hall. These partners are instrumental to the success of the CeVIVAS project.

## Supplementary Figures

**Figure S1.**
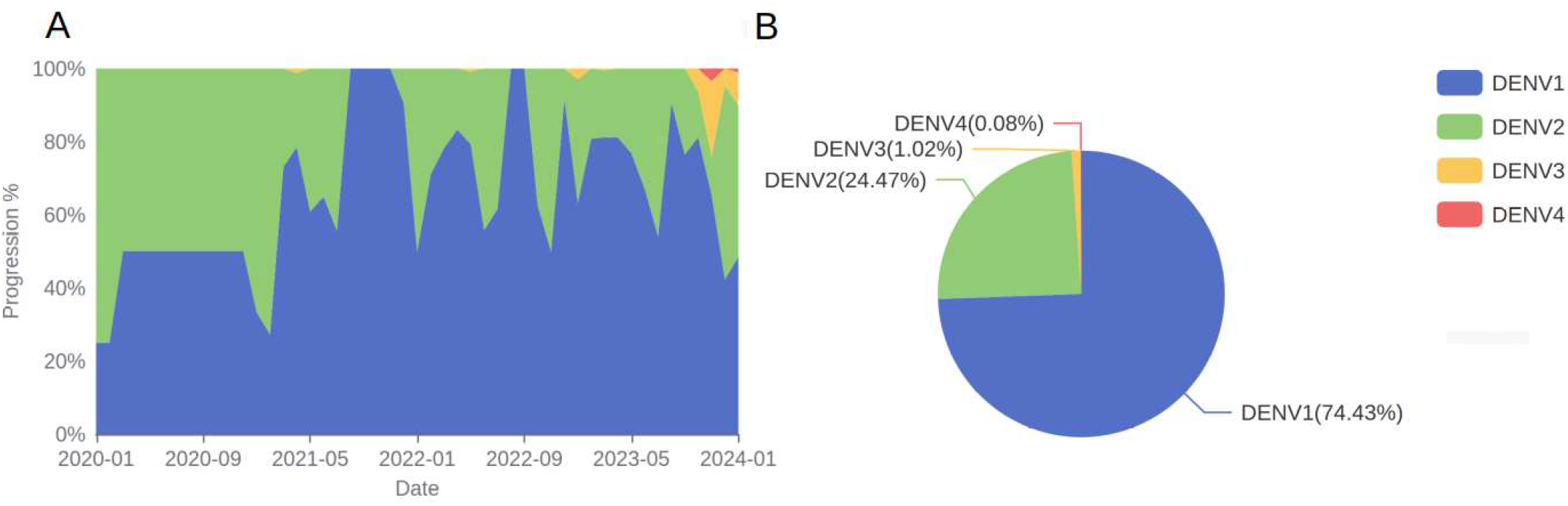
Distribution of DENV Genome Sequences across Serotypes in Brazil. **A.** Percentage distribution of DENV genome sequences by serotype, spanning from January 2020 to January 2024. **B**. Serotype prevalence of DENV based on genome sequencing data. Data sourced from GISAID as of March 15, 2024, encompassing 2,693 genome sequences.

**Figure S2.**
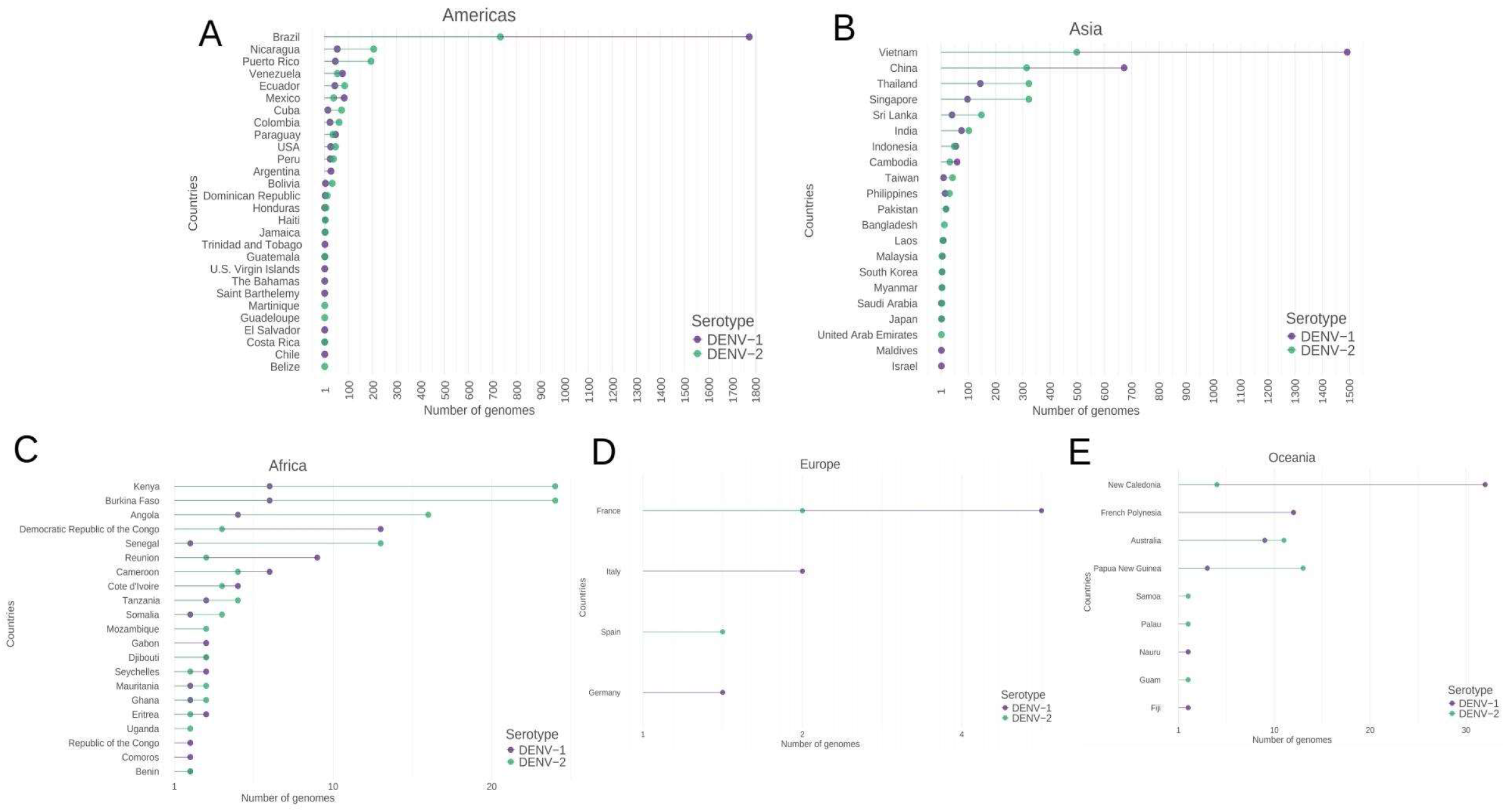
Geographical Distribution of DENV Genomes Employed for Lineage Assignment. The data is presented according to the continent and country of origin for each sequence.

## Notes

https://github.com/alex-ranieri/denvLineages

### Competing Interest Statement

The authors have declared no competing interest.

